# Acetyl-CoA metabolism drives epigenome change and contributes to carcinogenesis risk in fatty liver disease

**DOI:** 10.1101/2021.07.22.453333

**Authors:** Gabriella Assante, Sriram Chandrasekaran, Stanley Ng, Aikaterini Tourna, Carolina H. Chung, Kowsar A. Isse, Celine Filippi, Anil Dhawan, Mo Liu, Steven G. Rozen, Matthew Hoare, Peter Campbell, J. William. O. Ballard, Nigel Turner, Margaret J. Morris, Shilpa Chokshi, Neil A Youngson

## Abstract

The rate of nonalcoholic fatty liver disease (NAFLD)-associated hepatocellular carcinoma (HCC) is increasing worldwide, but the steps in precancerous hepatocytes which lead to HCC driver mutations are not well understood. Here we provide evidence that metabolically-driven histone hyperacetylation in steatotic hepatocytes can increase DNA damage to initiate carcinogenesis. Genome-wide histone acetylation is increased in steatotic livers of rodents fed high fructose or high fat diet. In vitro, steatosis relaxes chromatin and increases DNA damage marker γH2AX, which is reversed by inhibiting acetyl-CoA production. Steatosis-associated acetylation and γH2AX are enriched at gene clusters in telomere-proximal regions which contain HCC tumor suppressors in hepatocytes and human fatty livers. Regions of metabolically-driven epigenetic change also have increased levels of DNA mutation in non-cancerous tissue from NAFLD patients. Finally, genome-scale network modelling indicates that redox balance is a key contributor to this mechanism. Thus abnormal histone hyperacetylation is a potential initiating event in HCC carcinogenesis.

## Introduction

Hepatocellular carcinoma (HCC) is the most common type of primary liver cancer and the fourth most common cause of cancer-related death worldwide^1^. Its incidence is increasing across the globe and a major contributor to this increasing rate is the obesity epidemic and the concomitant increases in non-alcoholic fatty liver disease (NAFLD). At present NAFLD is estimated to affect around 25% of the world’s population^2^ and is the fastest growing cause of HCC in the USA, France and the UK^3^. Clinically NAFLD presents as a spectrum which progresses from simple steatosis to non-alcoholic steatohepatitis with accumulating inflammation and fibrosis culminating in cirrhosis and eventually liver failure. Whilst HCC risk is highest at the more severe stages, the risk remains even in fatty livers without cirrhosis^4,5^. HCC is a heterogeneous cancer and can display at least 6 subtypes each with a different combination of characteristic mutations and transcriptomes^6^. The most common genetic signature of HCC is mutation of the *Telomerase* (*TERT*) gene which is present in around 60% of cancers and is the earliest detectable of all the known mutations^7^. HCC-associated *TERT* mutations activate the gene, preventing telomere shortening which subverts cellular senescence and apoptosis programs thus setting cells on the path for immortalisation^7,8^. The underlying causes for *TERT* mutation can include viral (hepatitis B or C) insertion, point mutations which alter localised transcription factor binding, or larger rearrangements which duplicate the gene or translocate the regulatory sequences of a more highly expressed gene to the *TERT* locus^7^. However, the mutational forces which induce HCC-NAFLD are poorly understood. Oxidative stress is a strong candidate for generating mutations^9^ but the reasons why *TERT* mutations are so much more prevalent than other mutations at the early stages of carcinogenesis is unknown.

Diet and epigenetics are highly interlinked as the molecular substrates for epigenetic modifications are also products of intermediary metabolism^10^. This situation ensures that there is constant communication between the two processes, so that gene expression can be quickly modified to meet the energy demands of the cell. Perhaps the best understood example of the link between dietary macronutrients and epigenetics is histone acetylation. Acetyl coenzyme A (acetyl-CoA), is produced by the breakdown of carbohydrates^11,12^ and fats^13^ for cellular energy, as well as being the substrate for de novo lipogenesis (DNL) and ketone generation. It also provides the acetyl groups for histone acetylation, an epigenetic modification which can decondense, i.e. ‘open-up’ chromatin to facilitate DNA access for transcriptional or DNA repair proteins. *In vivo* measurements in rat have shown that absolute acetyl-CoA levels are increased in fatty liver^14^. In human NAFLD livers acetyl-CoA flux into DNL, and oxidation in the tricarboxylic acid (TCA) cycle, has been shown to be upregulated, while ketogenesis is reduced compared to healthy liver^15^.

The importance of epigenetic changes in cancer progression is well established, particularly in the context of changes which alter the transcription of oncogenes and tumor suppressor genes^16^. Mutations in epigenetic modifier proteins are commonly found in cancer, even as the initiating mutation^17,18^. The potential therapeutic opportunity of this relationship in the context of histone acetylation is currently being tested with several clinical trials of pharmacological inhibitors of the writers, histone acetyltransferases (HATs), and erasers, histone deacetylases (HDACs) in a variety of cancers, including HCC^19,20^. These therapies aim to reverse epigenetic dysregulation in partially or fully developed cancers, *after* cancer-promoting DNA mutations have occurred, rather than at the initial pre-mutational stages that promote carcinogenesis. This restricted use of epigenetic therapies is partly due to there being no previously reported examples of abnormal energy metabolism driving epigenetic dysregulation to initiate carcinogenesis. Considering the mechanistic links between metabolism, epigenetics and cancer, we hypothesised that perturbed acetyl-CoA metabolism and histone acetylation could be an unrecognised contributor to NAFLD pathology and HCC risk.

## Results

### Histone acetylation and chromatin state is altered in liver steatosis

To test our hypothesis, we first examined rodent and cell culture models of liver and hepatocyte steatosis. Using our established rodent models of NAFLD, we found in both the mouse model of fructose-diet induced liver steatosis (Fig 1A)^21^, and the rat model of high fat diet-induced steatosis (Fig 1B)^22^, an increased H4K16 acetylation compared to control diet animals. To dissect the mechanisms that link macronutrients and genome-wide epigenetic state, we treated immortalised human hepatocytes (IHH) with oleic acid for 4 days. This treatment quadrupled intracellular lipid content (Fig 1Ci-ii). We adapted a method for assessing genome-wide nuclear condensation in these cells^23^ and found that oleic acid treated cells had significantly decondensed chromatin compared to cells grown in control media (Fig 1D).

**Figure 1.**
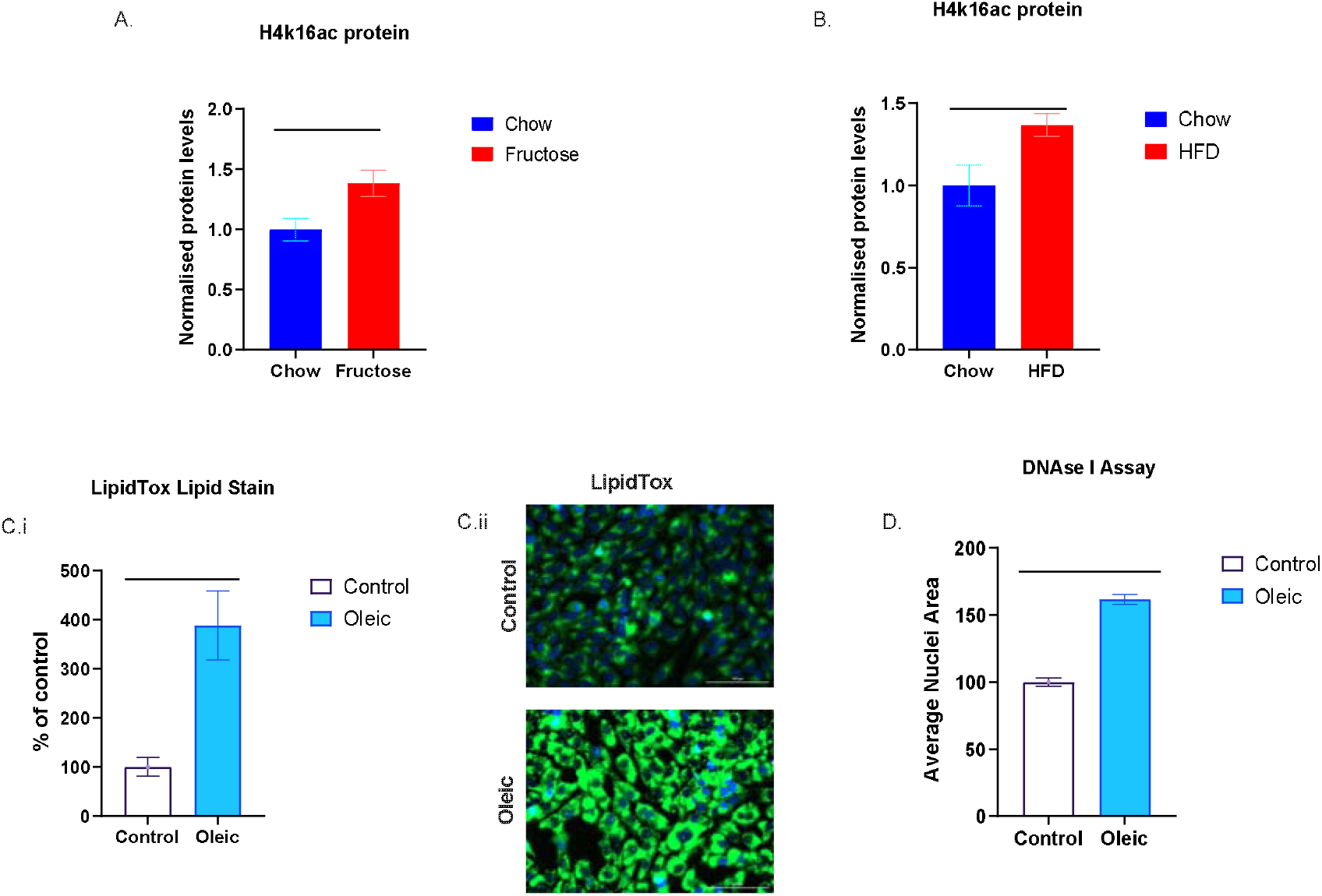
A. Histone acetylation levels in livers of control and high-fructose diet mice. B. Histone acetylation levels in livers of control and high-fat diet rats. Ci. Lipid content of IHH cells cultured in control or oleic-acid supplemented media. Cii. Representative fluorescence microscopy image of IHH cells cultured in control or oleic-acid supplemented media (blue nuclear stain and green lipid stain). D. Nuclear area measurement of IHH cells cultured in control or oleic acid supplemented media after DNAse I treatment. Error bars are ±SEM. Histone westerns all groups n=4, Lipidtox assay n=7/8, DNaseI Assay n=3 *P<0.05, ***P<0.001 in t-test.

### Hepatocyte steatosis is associated with increased ⍰H2AX which is reversed by inhibition of acetyl-CoA production

We assessed whether the global chromatin changes are associated with DNA damage by performing immunofluorescence for ⍰H2AX, a well-established marker of DNA damage^24,25^ Oleic acid treatment nearly doubled the percentage of IHH cells which had high levels of ⍰H2AX, compared to control media cells. However, this was reversed by a 4-hour pharmacological inhibition of any of the major sources of acetyl-CoA (Fig 2A, B). β-oxidation produces acetyl-CoA through the sequential breakdown of fatty acids. We reduced β-oxidation with the Carnitine Palmitoyltransferase-1 (CPT-1) inhibitor etomoxir. The ATP citrate lyase (ACLY) inhibitor (BMS-303141) prevents the production of acetyl-CoA from citrate, while acetyl CoA synthetase (ACSS2) inhibition prevents production of acetyl-CoA from acetate. Inhibition of all these sources of acetyl-CoA had the similar effect of reducing the number of cells with high ⍰H2AX to levels comparable with non-steatotic cells.

**Figure 2.**
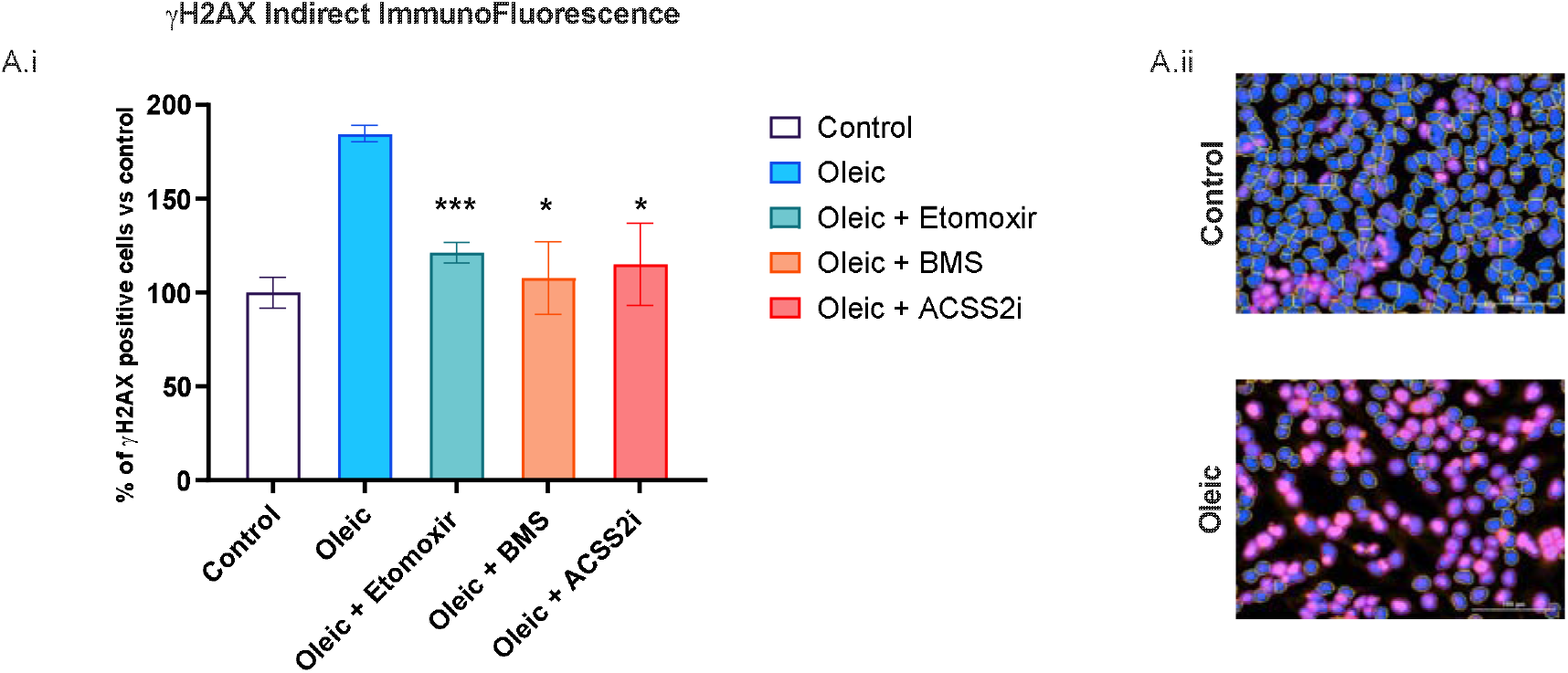
A. Average number of cells with high ⍰H2AX after oleic acid or oleic acid plus metabolic inhibitor drug treatments (% compared to control cells) B. Representative ⍰H2AX immunofluorescence images of control and oleic acid treated cells (blue nuclear stain, pink ⍰H2AX). Error bars are ±SEM. All groups n=6-8. *P<0.05, ***P<0.0001 in t-test of oleic vs oleic plus inhibitor groups

### Genomic regions with highest increases of ⍰H2AX levels in steatotic hepatocytes are clustered and contain genes commonly mutated in HCC

We next sought to identify which regions of the genome experience steatosis-associated increased ⍰H2AX. We performed chromatin-immunoprecipitation with anti-H4K16ac and anti-⍰H2AX antibodies and nextgeneration sequencing on the enriched DNA from control and steatotic IHH cells. The ⍰H2AX peak regions were compared between the control and steatotic cells. 77 of the 100 regions with biggest peak increases in steatosis compared to control cells were clustered on 7 chromosomes (5, 7, 11, 15, 19, Y and the mitochondrial genome). These clusters displayed colocalization of H4K16 acetylation and ⍰H2AX peaks (Supplementary Fig 1, Supplemental File 1). Two clusters stood out due to their telomere-proximal location and the genes within them. We searched The Cancer Genome Atlas for evidence that mutations have previously been found in these clusters in HCC. The Liver Hepatocellular Carcinoma Case Set (Project ID TCGA-LIHC) has information from 377 patients. 30 of the 32 protein coding genes in the chromosome 19 cluster of genes had identified mutations (most frequently at *BRSK1, ZNF135* and *EPS8L1* and *NLRP2*) in the patients, similarly 17 out of 21 genes at the chromosome 5 cluster had identified mutations (most frequently in *TERT, PLEGHG4B* and *IRX4*). The combination of the cluster-wide epigenetic change and identified mutations in multiple genes within these clusters supports the possibility that large chromosomal regions have increased mutation risk from metabolism-associated epigenetic change in steatosis.

### Genomic regions with highest increases of ⍰H2AX levels in steatotic hepatocytes have increased mutations in livers from NAFLD and alcohol-related liver disease patients

To investigate the potential clinical relevance of the epigenetic changes in steatotic IHH cells we compared the ChIP-seq data with genomic mutations in human liver biopsies from chronic liver disease patients. Previously we described the sites of DNA mutation in cirrhotic livers with differing underlying aetiologies^26^. Here we compared the number of single nucleotide variants (SNVs i.e. mutations) detected in these whole genome sequencing (WGS) datasets derived from primary human liver biopsies, with the ⍰H2AX peak regions in control media and oleic-acid-treated IHH cells. These comparisons confirmed that the regions with high ⍰H2AX levels in steatotic IHH cells were enriched in NAFLD and alcohol-related liver disease (ARLD) - associated SNVs (Fig 3B), whereas the regions with high ⍰H2AX levels in control media IHH cells were not (Fig 3A). Furthermore, a similar significant association was found between steatotic IHH ⍰H2AX peak regions and SNVs in the same cirrhotic livers of type 2 diabetes patients, but not with the control media ⍰H2AX peak regions (Supplementary Fig 2). These analyses suggest that the genomic regions which were identified to be sensitive to metabolism-associated epigenetic change in IHH cells are also regions of frequent mutations in diseased human livers.

**Figure 3.**
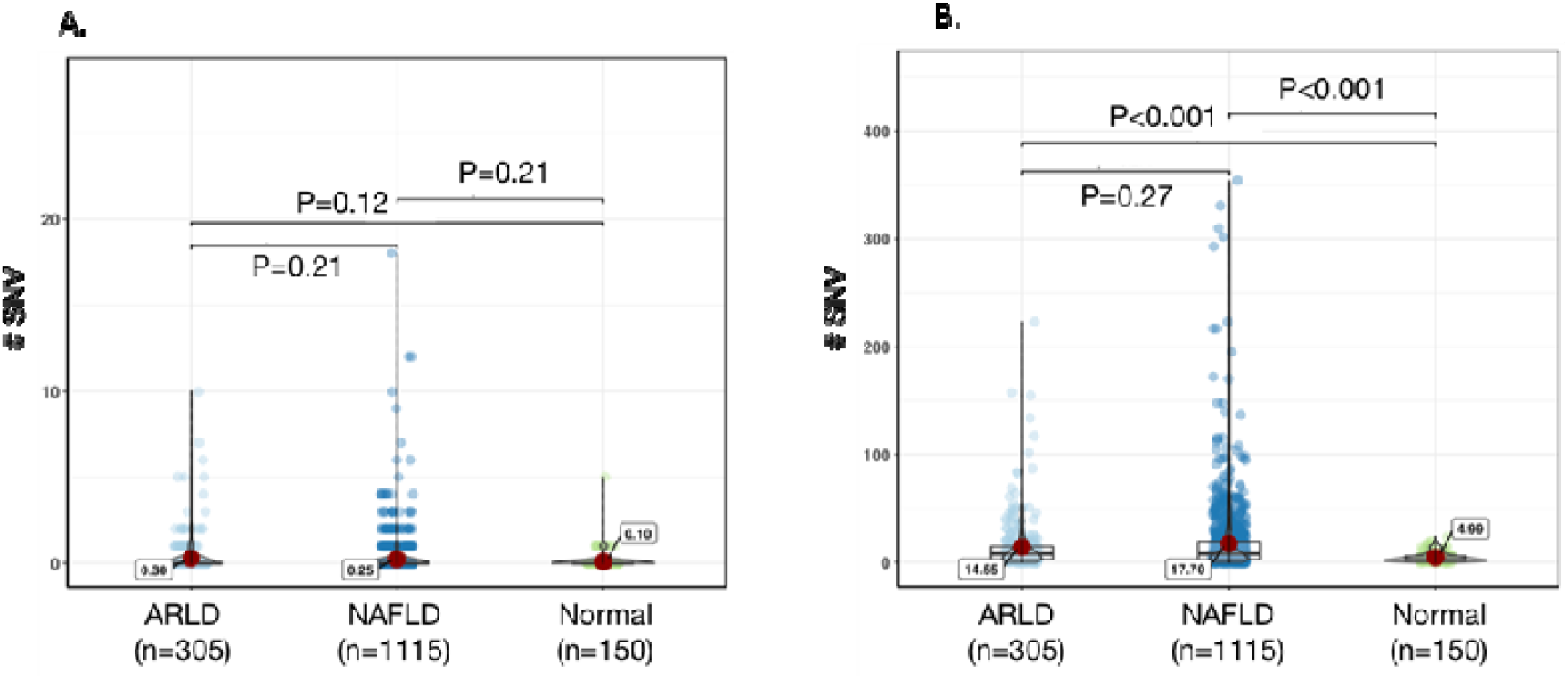
Violin plots showing a comparison of the number of single nucleotide variants (SNVs) in whole genome sequencing from laser capture microdissections of human liver from patients with normal liver (n=5), alcohol-related liver disease (ARDLD) (n=10) or Non-alcoholic fatty liver disease (NAFLD) (n=19), at A. control, and B. oleic acid peak regions. The median number of SNVs are annotated on the plots.

Subsequently we examined the mutational signatures that are present in the primary human liver biopsies WGS datasets at the regions which correspond to the ⍰H2AX peak regions. This indicated that the proportion of SNVs which are attributable to ROS-associated mutational processes are more pronounced in NALFD clones in the steatotic ⍰H2AX peak regions than control ⍰H2AX peak regions (Supplementary Fig 3). Indeed, the ROS-associated mutational signature had the greatest proportional increase in the ⍰H2AX peak regions between healthy and NAFLD livers.

### The *TERT* gene has high levels of histone acetylation and ⍰H2AX in steatosis which can be reversed with acetylation inhibitors

Mutations at the *TERT* gene are the earliest detected in HCC^7,8^. We validated the ChIP-seq results with ChIP-qPCR of the *TERT* promoter region compared to other tumour suppressor genes which are known to become mutated in later stages of hepatocellular carcinogenesis. This showed an approximate 40-fold increase in both H4K16ac and ⍰H2AX at the *TERT* promoter in steatotic IHH cells compared to control cells (Fig 4A). Near the PTEN gene, the increase in both histone modifications was more modest, while TP53 was even less changed (Fig 4A). Treatment of the cells for 4 hours with the ACLY inhibitor BMS or histone acetyltransferase inhibitor garcinol prior to ChIP-qPCR significantly reversed the steatosis-associated elevation in both H4K16ac and ⍰H2AX levels at the *TERT* promoter.

**Figure 4.**
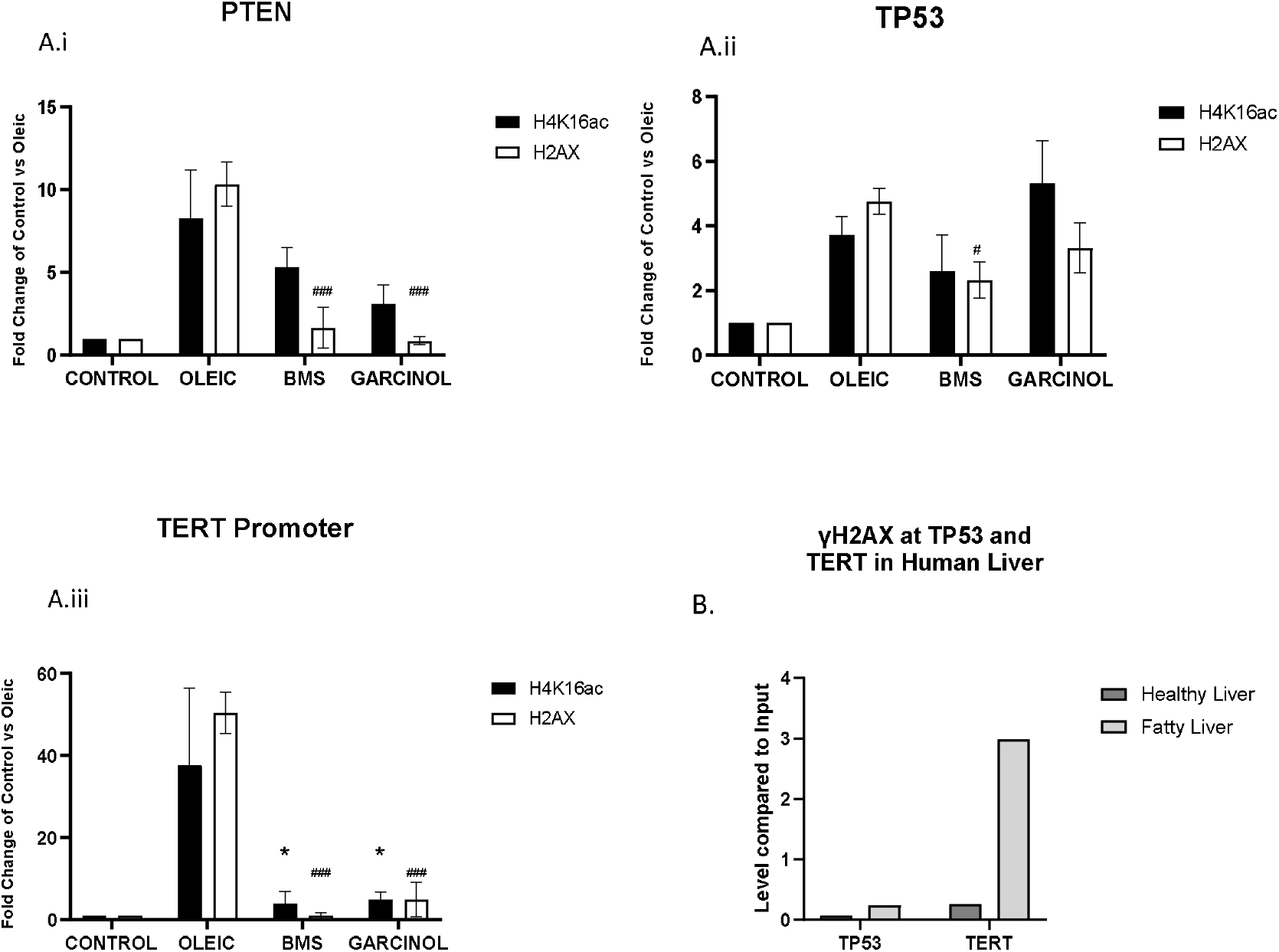
ChIP-qPCR analyses ⍰H2AX and H4K16ac levels at *PTEN* (Ai), *TP53* (Aii) and *TERT* promoter (Aiii) in IHH cells cultured in control media, oleic acid or oleic acid plus inhibitor drugs. Error bars are ±SEM. All treatments n=3. ^#,*^P<0.05, ^###^P<0.0001 in 1-way ANOVA, * H4K16ac and # ⍰H2AX. B. ChIP-qPCR analyses ⍰H2AX at *TP53* and *TERT* promoter in one human healthy liver and one human fatty liver (no statistical comparisons performed).

We were able to confirm the regional differences in ⍰H2AX levels at the *TERT* promoter and TP53 in human livers which had been rejected for transplantation (Fig 4B). A high levels of ⍰H2AX enrichment was only seen at TERT in fatty livers.

### Network modelling predicts metabolic consequences of oleic acid treatment and reversal by metabolic inhibitors

We used genome-scale metabolic network modeling to gain insight into how ⍰H2AX levels were increased by oleic acid and decreased by drugs which inhibit acetyl-CoA production^27 28^. We used the Recon1 metabolic model that contains the relationship between 3744 reactions, 2766 metabolites, 1496 metabolic genes, and 2004 metabolic enzymes curated from literature^29^. To create a hepatocyte-specific metabolic model, gene expression data from AML12 hepatocyte cells^13^ was overlaid onto the Recon1 network model (Methods). The model was further constrained using the nutrient conditions in culture and reaction fluxes were determined using an optimization approach that determines the metabolic flux state that satisfies constraints from gene expression, nutrient conditions, thermodynamics and stoichiometry^28^.

This analysis suggested that oleic acid treatment had numerous metabolic effects, impacting central carbon, fatty acid, amino acid and folate metabolic pathways. The model predicts that both the steatosis-associated increase of ⍰H2AX and its reversal is linked to redox metabolism. Many reactions which require the redox couples NAD+/NADH or NADP+/NADPH were altered in opposite directions in oleic acid treatment compared to the metabolic conditions induced by the inhibitors (Fig 5). Apart from the common alteration in redox metabolism, the three inhibitors were predicted to have distinct impact on the metabolic network. Both ACLY and CPT1 inhibition were predicted to increase folate metabolism, whereas ACSS2 inhibition did not.

**Figure 5.**
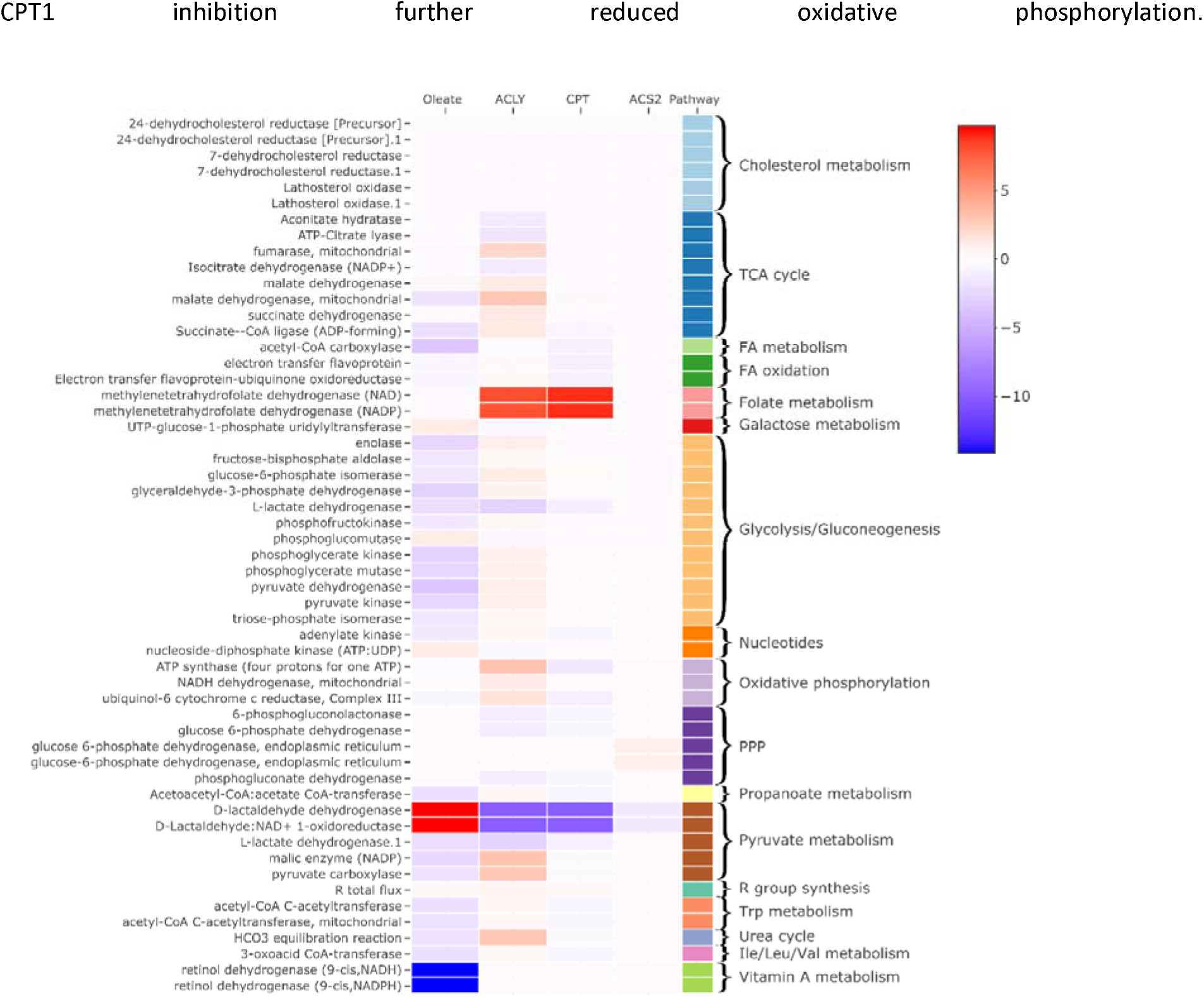
Genome scale metabolic modeling to simulate the impact of fatty acid treatment and metabolic enzyme inhibition on the metabolic network of hepatocytes. Heatmap of reaction differences between the oleic acid vs control medium and *ACLY, CPT, ACS* KO vs wild type control in oleic acid treated cells, p< 0.05 in any condition

## Discussion

In this study, we provide evidence that energy metabolism can direct epigenetic change and DNA damage across the genome, including to the earliest mutated gene in HCC. While acetylation changes and HDAC expression changes are known in HCC^20^ and NAFLD^30^, the relationship between histone acetylation and DNA mutations at known HCC-driver genes has not been reported. Our focus on non-cancerous, pre-mutational hepatocytes provides insight into the priming steps of an increasingly common cancer which has a five-year survival of 19%^31^ and is responsible for more than 700,000 attributable deaths per year worldwide^32^, and may explain why HCC risk increases in NAFLD patients from the early stages of steatosis. It is also to our knowledge, the first evidence for abnormal histone acetylation being the initiating epigenetic event in carcinogenesis.

Overall, our data support the concept that increased acetyl-CoA flux in steatotic hepatocytes leads to increased histone acetylation at certain genomic regions thus increasing their sensitivity to the prevailing oxidative stress. We observed that many of the genomic regions which had the highest increases in ⍰H2AX and H4K16ac in steatosis, also had relatively high levels (compared to the rest of the genome) in cells grown in control media. This suggests that these regions, such as the telomere-proximal cluster which contains *TERT*, may be predisposed for sensitivity^33^. There is an abundance of recent literature on the role of epigenetic modifications altering the expression of TERT in cancer via transcriptional and post-transcriptional mechanisms^34^, but the potential for the epigenetic states promoting mutagenesis has been hypothetical^35^. Despite the apparent risks for generating permanent mutations in pathologically important genes, the mutagenic metabolic environment can be rapidly reversed through alterations to energy and redox metabolism as shown with only 4 hours of treatment with various inhibitors. Further investigation is needed to confirm that this can be done safely *in vivo*.

The computational modelling of the metabolic changes induced by the different inhibitors of acetyl-CoA production suggest that oxidative stress is responsible for the ⍰H2AX decreases. This is supported by the presence of ROS-associated mutational signatures of the SNVs that are found at the genomic regions in fatty livers which correspond to the oleic-acid induced ⍰H2AX peaks. NAD/NADP-consuming enzymes in folate and pyruvate metabolism were predicted to be altered, and dysregulation of either process has been shown to influence oxidative stress in liver^36,37^. Telomeres are known to be particularly sensitive to damage due to oxidative stress^38^, further work is needed to determine whether subtelomeric regions also have particular sensitivity in steatotic hepatocytes. Additional mechanistic areas to investigate include evaluating the relative importance of oxidative stress, folate and pyruvate metabolism for epigenome change and DNA damage, and seeing how other NAFLD-promoting lipid species such as palmitic acid influence the process.

The worldwide rates of HCC are increasing and there is a switch from the incidence being driven by a relatively small population with viral hepatitis who have a high risk for HCC, to a large population with NAFLD who have a relatively low HCC risk^39^. Therefore, it is of interest to identify whether this new pathological mechanism is more pronounced in the small proportion of NAFLD patients who develop HCC. Our patient liver mutational analyses suggest that the same mechanism occurs in ARLD and T2D, which could potentially link increased HCC risk in the setting of steatohepatitis. Besides HCC, future work will aim to determine whether regionally increased histone acetylation can progress the severity of NAFLD to NASH and cirrhosis through dysregulation of genes involved in inflammation and fibrosis. Finally, several drugs undergoing preclinical and clinical trials for reducing liver steatosis and fibrosis in NAFLD will likely impact acetyl-CoA metabolism^40^ and/or histone acetylation^41–43^. For example, firsocostat is an acetyl-CoA synthetase inhibitor which blocks the use of acetyl-CoA for DNL.^44,46^. The effects of these drugs on histone acetylation and DNA damage should be evaluated, especially if long-term administration is being considered for chronic NAFLD.

## Methods

### Animals

All rodent experiments were approved by the University of New South Wales Animal Care and Ethics Committee. For the High Fat Diet treatment (HFD) male Sprague-Dawley rats from the Animal Research Centre (ARC, Perth, Australia) were housed two per cage under a 12:12 h light/dark cycle and *ad libitum* access to water and experimental diets. Three-week-old rats were split into two groups with equal average body weight (n = 14/14). Control rats were fed normal chow (energy: 11 kJ g-1, 12% fat, 21% protein, 65% carbohydrate; Gordon’s Stockfeeds, NSW, Australia) whilst the HFD group was provided with two commercial HFD pellets, SF03-020 (20 kJ g-1,43% fat, 17% protein, 40% carbohydrate; Specialty feeds, Glen Forest, WA, Australia) and SF01-025 (18.3 kJ g-1,44% fat, 17% protein, 39% carbohydrate; Specialty feeds), as well as normal chow. The rats were killed between 24 and 29 weeks of age. Animals were killed after anaesthesia induced by i.p. injection of 100 mg ketamine/kg body weight and 15 mg xylazine/kg body weight followed by decapitation. At killing the difference in body weight between the groups had increased so that the control group averaged ± s.d 535 ± 67 g (range: 451–631 g) and the HFD group 717 ± 67 g (range: 604–807 g), P = 6.0 × 10^-8^^22^.

For High Fructose diet (HF), ten-week old C57BL/6J mice were purchased from the Australian Resource Centre (Perth, Australia). Mice were maintained in a temperature-controlled room (22⍰°C⍰±⍰1⍰°C) with a 12-hour light/dark cycle and *ad libitum* access to water and experimental diets. After one week on a standard control “chow” diet (71% of calories from carbohydrate as wheat/starch, 8% calories from fat, 21% calories from protein, ~3kcal/g; Gordon’s Specialty Stock Feeds, NSW, Australia), mice were randomly allocated to remain on the chow diet (C) or to receive a home-made diet enriched in fructose (FR; 35% of calories from fructose, 35% calories from starch, 10% calories from fat, 20% calories from protein, ~3.1kcal/g) *ad libitum* for 8 weeks. At kill the control and fructose fed mice had no bodyweight difference but liver triacylglycerol levels in fructose mice were 225% the level of control mice p⍰<⍰0.001^21^.

### Western Blots (Immunoblotting)

Histones were extracted from approximately 30mg powdered liver on a Precellys (Sapphire Bioscience Australia) at 6⍰m/sec for 30⍰sec, in Triton Extraction Buffer (TEB: PBS containing 0.5% Triton X 100 (v/v), sodium butyrate 5mM) using the Abcam Acid Extraction Histone extraction protocol for western blot. Briefly, after lysis, samples are centrifuged at 6,500 x g for 10 min at 4°C to spin down the nuclei and supernatant discarded. Pellet resuspend in 0.2 N HCl and histones acid extracted overnight at 4°C on a rotator. The next day samples were centrifuged at 6,500 x g for 10 min at 4°C to pellet debris. The supernatant (which contains histones) was neutralised with 2M NaOH at 1/10 of supernatant volume. Equal amounts of protein lysate were electrophoresed through a 4–15% precast gel (Criterion TGX, Bio-Rad) for 45 min at 150 V in running buffer (25 mmol/l Tris base, 192 mmol/l glycine, and 1% SDS, pH 8.3). Proteins were transferred via a semi dry transfer process with a Trans Blot Turbo System (Bio-Rad) onto PVDF membranes (Bio-Rad). Membranes were blocked in 4% BSA in TBS-Tween for 1 h, then incubated overnight at 4°C with primary antibodies used at 1:2000 dilution; Histone H4K16ac pAb, (Active Motif 39167), total histone H3 (Abcam, abl79l). The membrane was subjected to three 10 min washes with TBS-Tween, and incubation with appropriate secondary antibody (CellSignaling) in 2% skim milk blocking solution in TBS-Tween at room temperature for 1 h, followed by three 10 min washes with TBS-Tween. For detecting bands, membranes were exposed to Clarity Western ECL Substrate (Bio-Rad) and visualized on a Bio-Rad ChemiDoc XRS. Membranes were stripped using Reblot Plus (10X) (Millipore) for 10 min at room temperature and were re-blocked prior to pan-H3 immunoblotting. Immunolabelled bands were quantitated using ImageJ 1.44p software.

### Cell culture

Immortalized Human Hepatocytes (IHH) were cultured under standard conditions by using control media DMEM/F-12 (Sigma) without phenol red, 10% FBS (Life Technologies, 10108-165), 1% PEN/STREP (Life Technologies 15140-122), 0.1% L-Glutamine (Life Technologies 25030-024), 0.02% Dexamethasone (Sigma D49025MG), 1pM Insulin Human Recombinant Zinc (Life Technologies 12585-014). For generating steatosis cells were supplemented oleic acid-albumin (Sigma-O3008) (300uM). After for 4 days of culture, confluent cells were treated with respectively with 200uM Etomoxir (Sigma-E1905), 50uM BMS 303141 (Sigma - SML0784), 5uM Garcinol (Thermo Fisher - 15716585) and 50uM ACCSi - CAS 508186-14-9 (Sigma - 5337560001) for 4 hours at 37°C.

### Lipid Droplets Staining

Intracellular lipid droplets staining was performed with HCS LipidTOX™ Green Neutral Lipid Stain (ThermoFisher). After removing the incubation medium, the cells were fixed with a solution of 3.7% of Formaldehyde (Sigma) supplemented with Hoechst 33342 (ThermoFisher) and incubated for 20 mins at room temperature in the dark. After aspirating the fixative solution, the cells were washed twice with PBS without Ca/Mg (Life Technologies). LipidTOX™ neutral lipid stain diluted in PBS was added to the cells and incubated for 1 hour at room temperature in the dark at a final concentration of 1x. Cell imaging acquisition was performed by using GFP 488nm and DAPI 355nm on Cytation 5 Cell Imaging MultiMode Reader (BioTek).

### In Situ DNAse I Sensitivity Assay: Chromatin State

CSK buffer was made by dissolving Pipes/KOH (Sigma), NaCl (Sigma), Sucrose (Sigma), EGTA (Sigma) and MgCl2 (Sigma) in H2O. Cover-slides were coated for 1 hour with 20% w/v of Fibronectin (Sigma) diluted in PBS (Life Technology) at room temperature. Next, cells were plated on the coated cover-slides, washed once with PBS and lysed in CSK buffer supplemented with 0.2% Triton X-100 (Sigma) and complete™ Protease Inhibitor Cocktail (Sigma/Roche) for 5 min at room temperature. Once the incubated solution was removed, the cells were washed with CSK buffer and then were incubated in CSK buffer supplemented with 0.1% Triton X-100 (Sigma), complete™ Protease Inhibitor Cocktail (Sigma/Roche) and 50 U/ml DNaseI (Sigma) for 20 min at room temperature. Cells were then washed with CSK buffer and the remaining DNA was stained using Hoechst 33342 (ThermoFisher) at a concentration of 5 ug/ml in CSK buffer supplemented with 125mM Ammonium Sulfate (Sigma) and complete™ Protease Inhibitor Cocktail (Sigma/Roche) for 5 min at room temperature. Cells were washed in CSK buffer and fixed in 100% Methanol (Sigma) for 5 min at −20°C. The methanol was removed and the CSK buffer was added before imaging nucleus areaacquisition by using DAPI 355nm on Cytation 5 Cell Imaging Multi-Mode Reader (BioTek). The Nucleus area of cells was acquired by a mask function created on the Microscope Software.

### Indirect ⍰H2AX Immuno-Fluorescence

Control and Steatotic cells were washed twice in PBS (Life Technology) after 4 hour drug treatment. Next, cells were fixed with 4% solution of Formaldehyde (Sigma) for 15 min at room temperature and washed twice in PBS. Then, cells were permeabilized with 0.25% Triton X-100 (Sigma) for 10 min at room temperature, washed twice in PBS and incubated with 3% BSA (Sigma) for 1 hour at room temperature. Cells were first incubated with Anti-gamma H2A.X (phospho S139) antibody - ChIP Grade (Abcam) for 1 hour at 37°C and then incubated with fluorochrome-conjugated secondary antibody (Sigma SAB4600400) for 1 hour at 37°C in the dark. Cells were then stained with 1ug/mL Hoechst 33342 (ThermoFisher) for 10 min at room temperature and the imaging acquisition was performed by using DAPI 355nm and RFP 558nm on Cytation 5 Cell Imaging Multi-Mode Reader (BioTek). The percentage of positive cells was calculated with a mask function created on the Microscope Software.

### ChIP and Next Generation Sequencing

Chromatin was isolated from 20 million cells using the Magna ChIP A/G Kit (One-day chromatin Immunoprecipitation Kits, Merck) according to manufacturer’s instructions. Chromatin was sonicated with a Vibra-Cell VCX750 probe sonicator (Sonics) for 3 pulses of 15s separated by 30s intervals at 30% output. 50ul of chromatin was immunoprecipitated with 5ug antibody specific for Anti-gamma H2A.X (phospho S139) antibody - ChIP Grade (Abcam) or with 5ug antibody specific for histone H4k16ac (ActiveMotif). 100-250ng of isolated DNA (average size 500bp) was sent to Genewiz for sequencing in an Illumina HiSeq 2×150bp. Sequencing yield was 29-37Mb per sample. Sequence analysis was also performed by Genewiz. Sequence reads were trimmed to remove possible adapter sequences and nucleotides with poor quality at 3 end (error rate > 0.01) using CLC Genomics Server 9.0. Trimmed data was then aligned to reference genome for human genome hg38. During the mapping, only specific alignment was allowed. 163-198M reads were aligned (all >95%). Peak analysis for each sample was done using the Histone model algorithm. As a result, a list of peaks (p < 0.05) was obtained from each treatment sample. Detected peak sequences were extracted and peak coverages were calculated. Total peak number from control media samples were 2815 and 1291 from the H4K16ac and ⍰H2AX ChIP-seqs, respectively. Total peak number from oleic acid treated samples were 650 and 5912 from the H4K16ac and ⍰H2AX ChIP-seqs, respectively.

### ChIP and quantitative PCR

The King’s College Hospital, London, ethics number for hepatocyte biology held by Professor Dhawan and Dr Filippi is LREC protocol 1998-0249. Chromatin was isolated from 20 million IHH cells or 40mg of of human fatty liver from donor patients (using the EpiQuik Chromatin Immunoprecipitation Kit (Epigentek) according to manufacturer’s instructions for cells or tissues. Sonication on isolated chromatin was performed as for the ChIP-seq for cells but with pulses increased to 4 for tissue. Real-time PCRs were performed in triplicate with primers for *TERT* promoter, PTEN and TP53 by using Luna^®^ Universal qPCR Master Mix (NEB - M0003) as SybrGreen Probe on an Ariamx Real-Time PCR System (Agilent). Primer sequences for qPCR were as follows: PTEN ChIPqPCR Forward *5’-GAGTCGCCTGTCACCATTTC-3’* PTEN ChIPqPCR *Reverse 5’-GCGCACGGGAGGTTTAAAA-3’*; TERTp ChIPqPCR *Forward 5’-GGATTCGCGGGCACAG*ACS’, TERTp ChIPqPCR *Reverse 5’-AGCGCTGCCTGAAACTCG-3’*; TP53 ChIPqPCR *Forward 5’-GTACCACCATCCACTACAACTACATGT-3’*, TP53 ChIPqPCR *Reverse 5’-GGCTCCTGACCTGGAGTCTTC-3’*.

### Statistical analysis for wet-lab work

Results are expressed as mean ± SEM. Data were analyzed using Student’s t-test or one-way ANOVA, followed by *post hoc* LSD tests using Graphpad Prism.

### Mutation burden comparisons

The ethics number under which the samples were acquired for mutational burden comparisons and mutational signature extraction was Cambridge (16/NI/0196). Comparisons between the number of SNVs detected in a whole genome sequencing (WGS) dataset derived from primary human liver biopsies (Ng et al., in revision at Nature, 2021) between various clinically annotated conditions was performed in the R statistical programming using the ggstatsplot package^47^. For all comparisons, the default non-parametric test was used as defined by the software, while Benjamini-Hochberg method was enabled to correct p-values for multiple hypothesis testing. The normal liver donors were 5 colorectal cancer patients with average Kleiner Fibrosis Scores of 0.8 and average BMI of 26.6, 19 NAFLD patients had an average Kleiner Fibrosis Score of 3.2 and average BMI of 30.8, 10 ARLD patients had an average Kleiner Fibrosis Score of 3.9 and average BMI of 26.1.

### Mutational signature extraction

Single base substitution signatures in 96-trinucleotide contexts was extracted using the mSigHdp (v1.1.2) and hdpx (v0.3.0) package in R. 50,000 burn-in iterations were used, while setting the following hyperparameters and posterior sampling variables: post.n = 200, post.space = 100, gamma.alpha = 1, gamma.beta = 20. All other parameters were set to defaults. Cosine similarity was calculated to compare the extracted mutational signatures to the compendium of single base substitution COSMIC signatures^48^.

### Computational metabolic modelling

Genome-scale metabolic models have been widely used to predict the metabolic behaviour of various mammalian cell types using transcriptomics data^49,50^. Gene expression data from mouse hepatocyte AML12 cells treated with fatty acids (Octanoate) from McDonnel et al^13^ was used as input to derive reaction flux through the human genome-scale metabolic model (RECON1)^29^. The human orthologs of the lists of up- and down-regulated mouse genes were overlaid onto the RECON1 model based on gene-protein-reaction annotations in the model. Reaction fluxes that best fit the expression data while satisfying stoichiometric and thermodynamic constraints were determined using a modeling approach detailed in Shen et al^27,28^ This approach maximizes flux through reactions that are up-regulated while minimizing flux through those reactions that are down-regulated using linear optimization. The exchange reactions for nutrients (i.e. glucose, amino acids, fatty acids, vitamins and minerals) in the metabolic model were constrained based on media composition used in this study (DMEM-F12 media with 500uM oleic acid). For visualization, a z-score transformation was performed on the flux difference between control and treatment groups, and the significance of the difference was determined using a paired t-test. Reactions showing significant changes in flux (p-value < 0.05) were visualized in the heatmap (excluding transport-, exchange-, and pseudo-reactions).

### Data Availability

ChIP-seq data are available in the NCBI Short Reads Archive, BioProject accession number PRJNA741105 at http://www.ncbi.nlm.nih.gov/bioproject/741105.

## Supporting information

Supplemental File 1 - ChIP-seq peak regions

## Acknowledgements

This study was supported by core funding from the Foundation for Liver Research, the Australian Research Centre Discovery Project Grant (DP190102555) and MOH-000032/MOH-CIRG18may-0004 and the Singapore Ministry of Health via the Duke-NUS Signature Research Programmes (SGR).

## Author contributions

G.A. performed cell culture experiments. S. Chandrasekaran and C.H.C. performed computational modelling; S.N. performed diseased liver SNV comparison to ChIP-seq regions under supervision of P.C.; S.N. also performed mutation signature analysis with M.L. and S.R., K.A.I. performed TCGA database analysis; A.T. performed in-silico transcriptional analyses; C.P. and A.D. provided transplant-rejected human liver; experimental concept developed by J.W.O.B., N.T., M.J.M, S. Chandrasekaran and N.A.Y.; N.T. and M.J.M also provided rodent models; S. Chokshi guided HCC research and contributed to cell culture experimental design; N.A.Y. also undertook rodent molecular analyses, designed cell culture experiments and drafted manuscript.

---

**Supplementary Figure 1.**
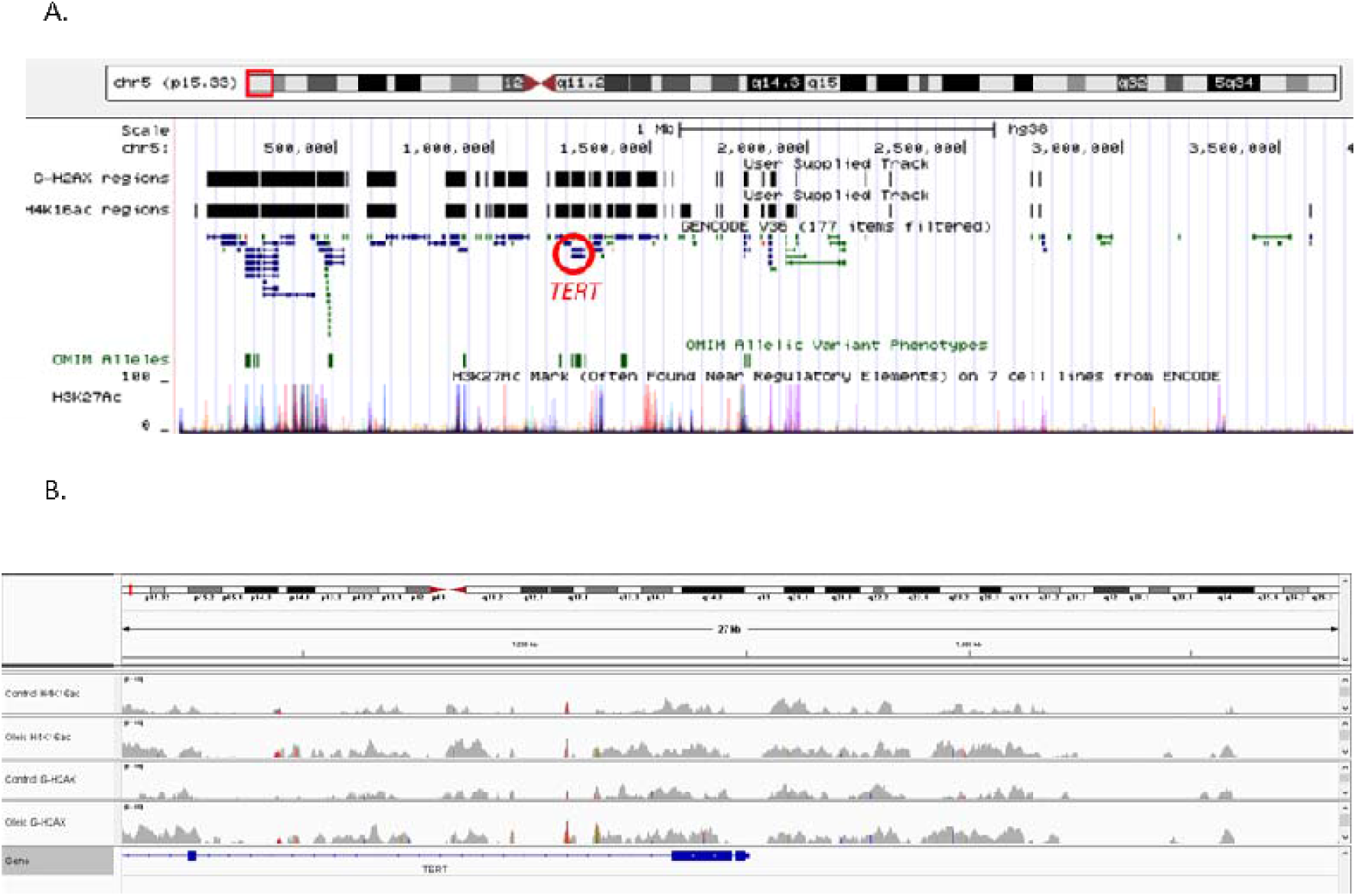
A. UCSC Genome Browser of the chromosome 5 telomere-proximal gene cluster showing location of genes and ENCODE H3K27ac regions relative to ⍰H2AX and H4K16ac ChIP-seq peak regions. B. IGV Genome Browser snapshot of the *TERT* promoter regions displaying aligned sequence reads from ChIP-seq files of the 4 samples top to bottom, control media cells H4K16ac, oleic acid treated cells H4K16ac, control media cells ⍰H2AX, oleic acid treated cells ⍰H2AX.

**Supplementary Figure 2.**
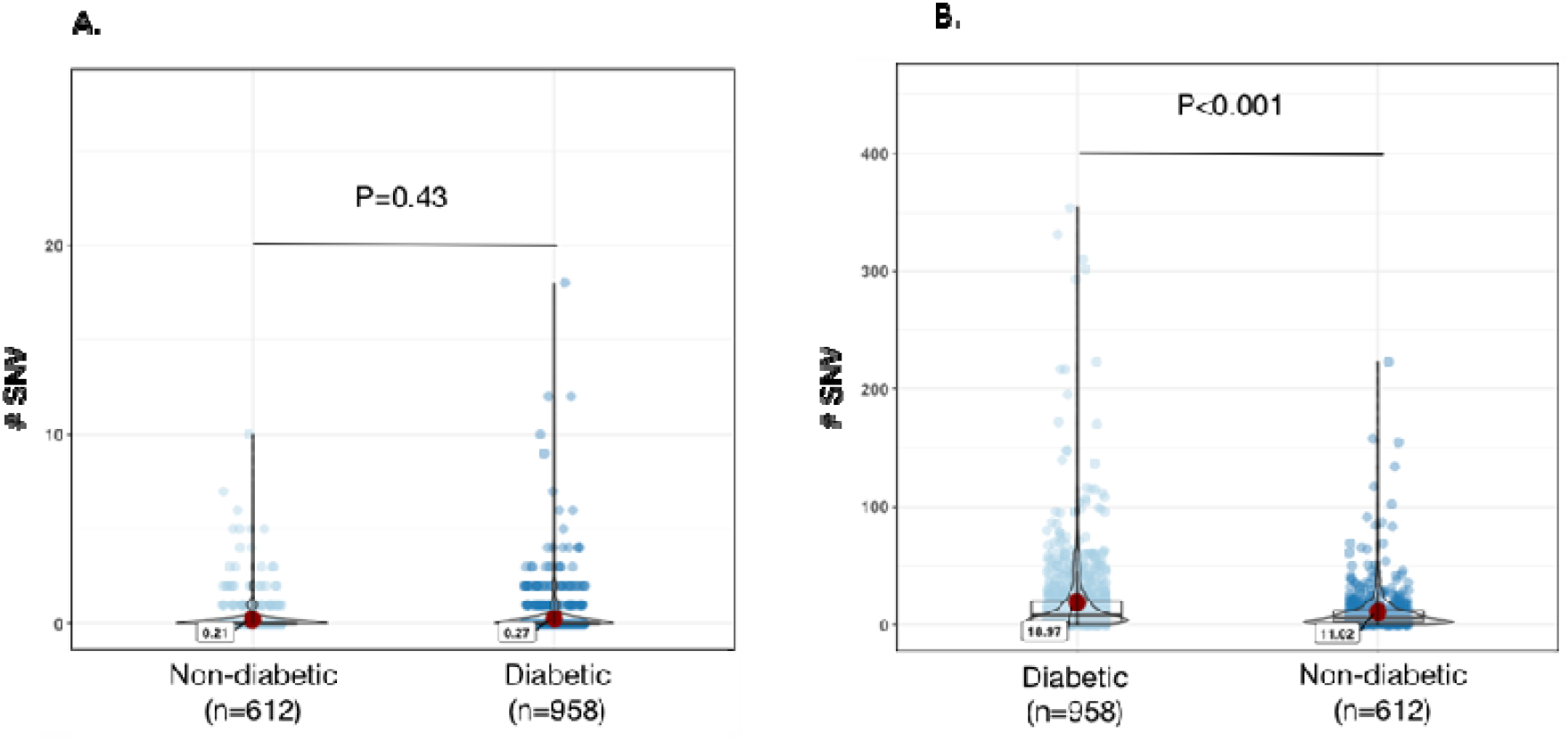
Violin plots showing a comparison of the number of single nucleotide variants (SNVs) in individual genetic clones from donors with versus without diabetes, at A. control, and B. oleic acid peak regions. The median number of SNVs are annotated on the plots. Genetic clones from 20 non-diabetic patients, and 14 diabetics.

**Supplementary Figure 3A.**
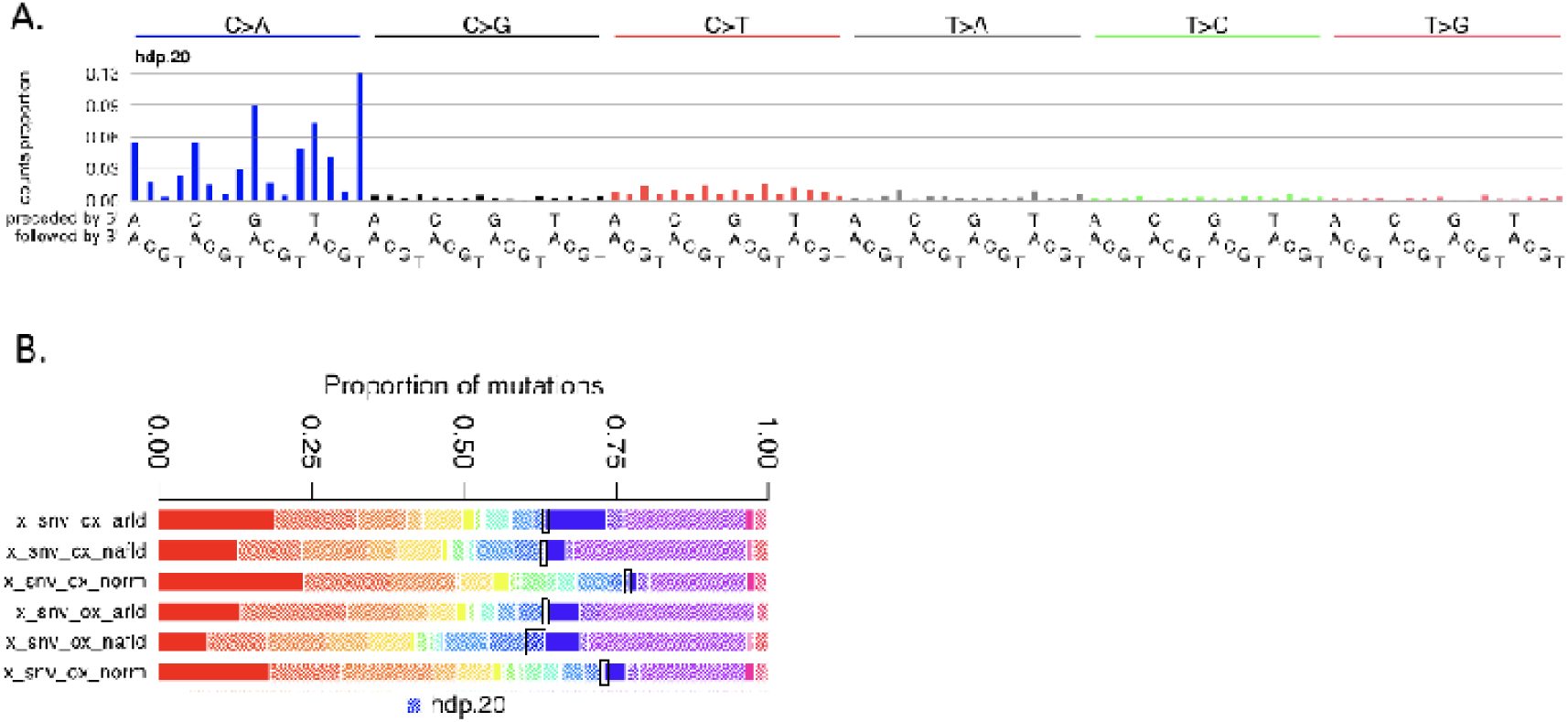
A mutational signature (hdp.20) was detected with 97% cosine similarity to SBS18, indicating that a proportion of SNVs is attributable to a ROS associated mutational process and was most pronounced in NALFD clones at oleic acid peak regions B. Bold black outlines highlight the relevant regions on the stacked bar plots corresponding to the ROS associated mutational signature. CX indicates control media cells ⍰H2AX ChIP-seq peak regions, OX indicates oleic acid treated cell ⍰H2AX ChIP-seq peak regions.

